# Mandibular morphology, task specialization, and bite mechanics in *Pheidole* ants (Hymenoptera: Formicidae)

**DOI:** 10.1101/2020.11.16.385393

**Authors:** C.L. Klunk, M.A. Argenta, A. Casadei-Ferreira, E.P. Economo, M.R. Pie

## Abstract

The remarkable ecological and evolutionary success of ants was associated with the evolution of reproductive division of labor, in which sterile workers perform most colony tasks whereas reproductives become specialized in reproduction. In some lineages, the worker force became further subdivided into morphologically distinct subcastes (e.g. minor vs. major workers), allowing for the differential performance of particular roles in the colony. However, the functional and ecological significance of morphological differences between subcastes is not well understood. Here, we applied Finite Element Analysis (FEA) to explore the functional differences between major and minor ant worker mandibles. Analyses were carried out on mandibles of two *Pheidole* species, a dimorphic ant genus. In particular, we test whether major mandibles evolved to minimize stress when compared to minors using combinations of tooth and masticatory margin bites under strike and pressure conditions. Majors performed better in pressure conditions yet, contrary to our expectations, minors performed better in strike bite scenarios. Moreover, we demonstrate that even small morphological differences in ant mandibles might lead to substantial differences in biomechanical responses to bite loading. These results also underscore the potential of FEA to uncover biomechanical consequences of morphological differences within and between ant worker castes.

## Introduction

The evolution of highly complex societies in ants was associated with the evolution of reproductive division of labor, in which sterile workers perform most quotidian colony tasks, whereas reproductives become specialized in colony reproduction (Wilson 1971, Hölldobler & Wilson 1990). These changes were accompanied by substantial morphological differences among reproductives and workers, with the latter giving up reproduction and dispersal capacities to allow for morphological adaptations that improve the ergonomic efficiency of the colony as a whole. In some ant lineages, the worker force became further subdivided into morphologically distinct subcastes (e.g. minor vs. major workers), and such differences are thought to allow for the differential performance of particular roles in the colony, such as seed milling and colony defense (Wilson 1953; Oster & Wilson, 1978). In ants, polymorphism evolved in several lineages, and its role to facilitate task specialization is widely recognized (Wilson 1953; Oster & Wilson, 1978; Wills et al., 2018), and the genetic (Gadagkar 1997; Huang et al., 2013), ecological (Powell & Franks, 2005; 2006; Powell 2009) and developmental (Wheeler 1991) determinants associated with the occurrence of worker polymorphism have been explored in several ant lineages (Wills et al., 2018).

The genus *Pheidole* shows an interesting pattern among its almost 1,200 known species (Bolton 2020), the development of a dimorphic worker subcaste, represented by major and minor workers, where majors have a disproportionally larger head (Wilson 1953; 2003). *Pheidole* species are distributed worldwide, but most of its diversity and abundance is concentrated in the tropics (Economo et al., 2015a; 2019). Although *Pheidole* species are typically considered diet generalists (Wilson 2003), some species might show degree of food preferences (Rosumek 2017). Of all their food items, feeding on seeds evolved many times in the genus and has been indicated as an important factor to explain the lineage diversification due to behavioral and morphological adaptations related to seed harvesting and processing (Moreau 2008). Since majors are specialized in tasks as colony defense and food processing (Wilson 1984; 2003), their larger heads could be a consequence of evolutionary pressures towards the specialization to those tasks (Pie & Traniello, 2007), although evidence gathered so far has been mixed (e.g. Holley et al., 2016).

Understanding the main trends in the morphological evolution of *Pheidole* received considerable attention in the past decade, with different approaches being employed to understand the evolution of a wide variety of structures, showing contrasting results to the relative contributions of size and shape to the morphological diversity of the genus (Pie & Traniello, 2007; Pie & Tschá, 2013; Sarnat et al., 2017; Friedman et al., 2019; 2020), yet little is known about the evolution of mandibular morphology in *Pheidole*. Ants have a typical pterygote mandible with two articulations with the head (Snodgrass 1935), the dorsal and ventral mandibular joints (Richter et al., 2019). The mandible is the main structure used to interact with the environment (e.g. biting, carrying, excavating, cutting, fighting) (Wheeler 1910). Mandibular movement is led by two muscles, the *craniomandibularis internus* (*mci*), whose contraction close the mandibles, and the *craniomandibularis externus* (*mce*), responsible for the opening process (Snodgrass 1935; Richter et al., 2019). The *mci* fibers attach to the mandible through a mandibular cuticular projection called apodeme (Paul & Gronenberg, 1999). The angle of attachment to the apodeme, combined to the sarcomeres length, are directly related to the velocity and force of the mandibular movement (Paul & Gronenberg, 1999), so that *mci* is considered the key to the versatility of ant mandibles (Gronenberg et al., 1997; Paul 2001), being much more developed than the *mce* (Paul 2001). In *Pheidole* majors the *mci* is remarkably large, where its increase in size in relation to minors happens at the expense of the glandular, digestive, and nervous system in the head (Lillico-Ouachour et al., 2018). Fibers of the *mci* continue to develop even for days after the adult emergence in both subcastes, and this characteristic correlates to behavioral development in workers (Muscedere et al., 2011).

Regardless of the importance of mandibles to many aspects of ant life history, little is known about how the morphological variation between species or worker subcastes relates to bite loads, except for one specialized snap-jaw species (Larabee et al., 2018). Worker polymorphism can lead to behavioral specialization, mainly through variation in mandible morphology (Silva et al., 2016), but biomechanical approaches to directly assess this relationship in ants are scarce (Larabee et al., 2018). To understand how mandible morphology relates to the biomechanical demands of bite, it is important to employ approaches that allow for the direct assessment of bite loading conditions. Finite Element Analysis (FEA) is a numerical method that approximates the mechanical simulation of loading conditions in structures of interest. By applying loads and defining the boundary conditions (movement restrictions) on the structure, FEA estimates the mechanical response, i.e., how stress flows along the structure according to its shape (Azevedo 2003; Rayfield 2007). By employing FEA, one might assess how variation in mandibular morphology among ant species as well as between subcastes translates into the capacity of mandibles to deal with bite loading demands (Larabee et al., 2018), as also explored for the evolution of mandible form in dragonflies (Blanke et al., 2017) and stag beetles (Goyens et al., 2015; 2016).

To improve our understanding of morphological evolution in *Pheidole* species, and the role of morphological differentiation to improve task specialization in polymorphic ants, we simulate several bite scenarios *in silico* by applying FEA (Azevedo 2003; Rayfield 2007) on 3D models of minor and major mandibles of two *Pheidole* species. We hypothesize that major mandibles are better able to mitigate stress than minors’, given their greater robustness. Alternatively, if each worker type has mandibles optimized to perform different tasks, majors and minors could perform better in distinct biting scenarios. Differences between species are expected between the more distinct mandibles of majors, which can suggest changes in the capacity to deal with hard food items, given the specialized roles of those workers (Wilson 1984; 2003). Alternatively, differences between species in minor worker mandibles will suggest that even small morphological distinctions can lead to biomechanical idiosyncrasies.

## Methods

### Studied species

Colonies of *Pheidole hetschkoi* Emery and *P*. cf. *lucretii* were collected in an urban fragment of Atlantic Forest in Curitiba, Paraná, Brazil (25°26’45.9”S 049°13’55.5”W). *Pheidole hetschkoi* is commonly found nesting fallen twigs; foraging in the species is based on the recruitment of tens of workers to food sources (rarely majors), and colonies accumulate seeds inside their nests (author’s pers. obs.). On the other hand, *P*. cf. *lucretii* nests directly on the ground and might recruit up to hundreds of workers to food sources, including dozens of majors, but were never recorded collecting or consuming seeds (author’s pers. obs). Morphologically, majors of *P. hetschkoi* are more sturdy, with larger heads (Fig.1b), with more robust mandibles than *P*. cf. *lucretii* majors, which have also smaller heads and are more slender in general (Fig.1a). Minors differ little between species in terms of mandible shape and general morphology (Fig. 1c and d). Voucher specimens are deposited at the Entomological Collection Padre Jesus Santiago Moure, Department of Zoology, Federal University of Paraná, Brazil.

**Fig. 1.**
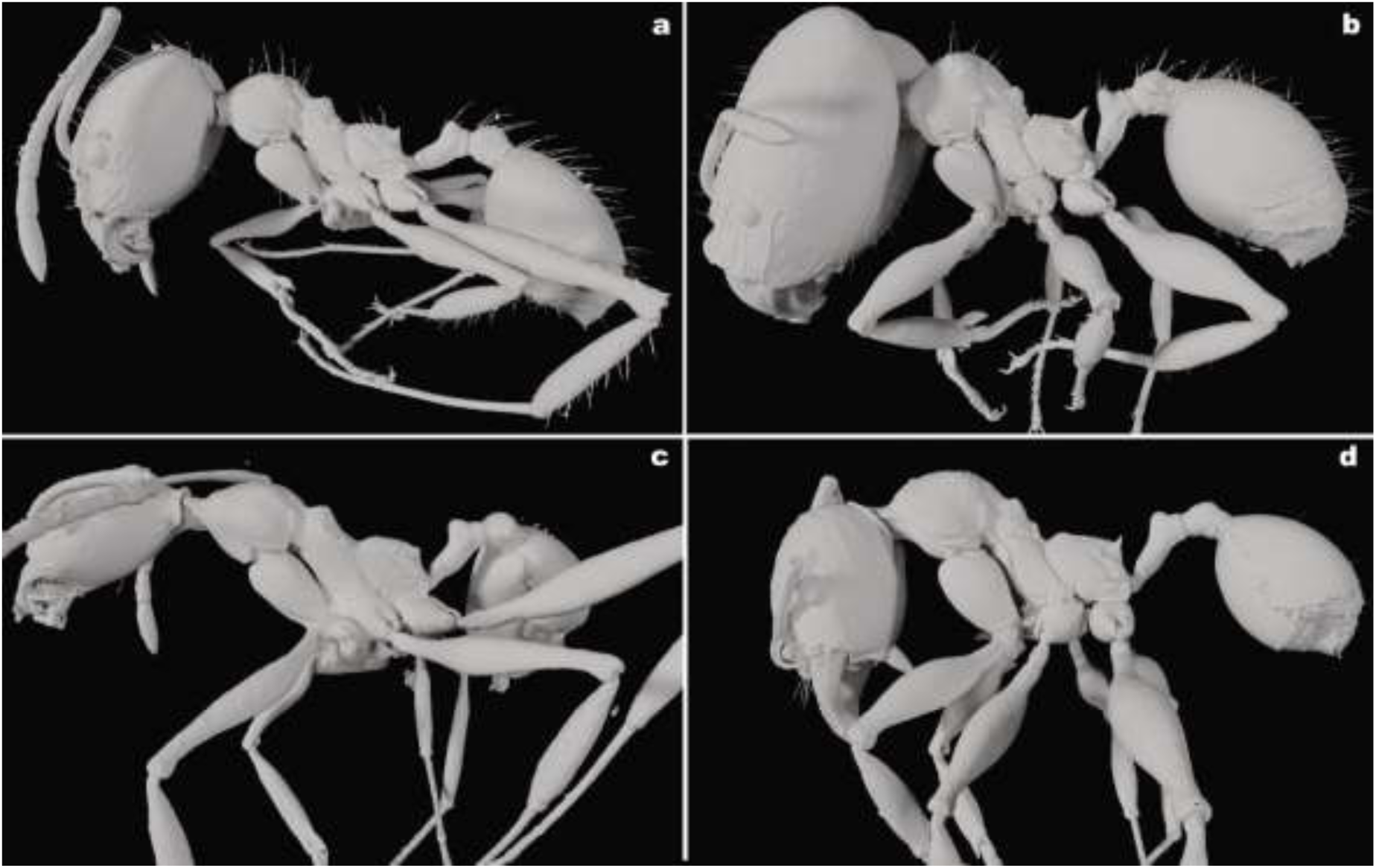
*Pheidole* workers whole body 3D models in lateral view. (a) *P*. cf. *lucretii* major; (b) *P hetschkoi* major; (c) *P*. cf. *lucretii* minor; (d) *P hetschkoi* minor.

### CT scanning and image processing

Workers scans were generated in a ZEISS Xradia 510 Versa X-ray microCT facility, using the software ZEISS Scout and Scan Control System, in the Biodiversity & Biocomplexity Unit, Okinawa Institute of Science and Technology Graduate University (OIST), Japan. Exposure time of each specimen varied from one to five seconds, under an “Air” filter and 4x objective. The voltage was set between 30 and 50keV, from 4 to 5W of power, under a “normal” field mode and intensity levels of 15,000 and 17,000 across the whole specimen. Scan time varied from 27 to 30 minutes, generating 801 projections from full 360-degree rotations. Model reconstruction was done on XMReconstructor, and mandibles segmentation was carried in ITK-snap 3.8.0 (Yushkevich et al., 2006). For mesh generation and simplification, we used the software MeshLab (Cignoni et al., 2008), and to generate the 3D mandible solid for FEA simulations we used the software Fusion 360 (AUTODESK). Mandibles 3D solid models are available on the supplementary material.

### FEA simulations

To quantify the mechanical response of a structure to external loading, FEA requires the discretization of the structure into small parts, resulting in the finite element mesh composed of elements of pre-defined shape and a specific number of points, called nodes, used to solve the equations (Azevedo 2003, Marcé-Nogué et al., 2015). Displacements on nodes are calculated to estimate stress and strain, based on the structure material properties and shape (Rayfield 2007). We used 10-node tetrahedral elements (C3D10) to generate the finite element mesh. The number of elements varied for each model, as well as the size of each element between subcastes, to adapt meshes to each morphology (Table SI1).

We performed linear static simulations of four distinct biting scenarios for each species and subcastes, divided into two categories, namely strike and pressure, which reflect different aspects of mandible movement in terms of force and velocity. In strike scenarios, a condition associated with faster movements, we define the mandible articulations with the head (dorsal - *dma* and ventral - *vma*) as the constrained regions, applying static load on the apical tooth or the masticatory margin (Figure 2a, b). In pressure scenarios, associated with slower mandible movements but powerful bites, in addition to the mandibular joints, we also constrained the apical tooth or the masticatory margin and applied the load to the region of *mci* insertion, following the direction of contraction (Figure 2a, c) to simulate the use of mandibles for food compression. We constrained nodal displacement in x, y, and z directions and apply a 1 N load uniformly distributed among nodes in all simulations. We modeled the mandible cuticle as an isotropic and linearly elastic material, setting Young’s modulus as 2.75GPa and the Poisson’s ratio as 0.3, based on measures from the cuticle of ant mandibles available in the literature (Brito et al., 2017). The only source of variation for each biting simulation between species and workers was the morphology of the mandibles. We present FEA stress results based on Tresca failure criterion, more suitable for brittle fracture, which determine an equivalent stress value under which the material will possibly fail when subjected to combined load (Özkaya et al., 2017). We used Abaqus 6 (Dassault Systèmes) to run the FEA simulations.

**Fig. 2:**
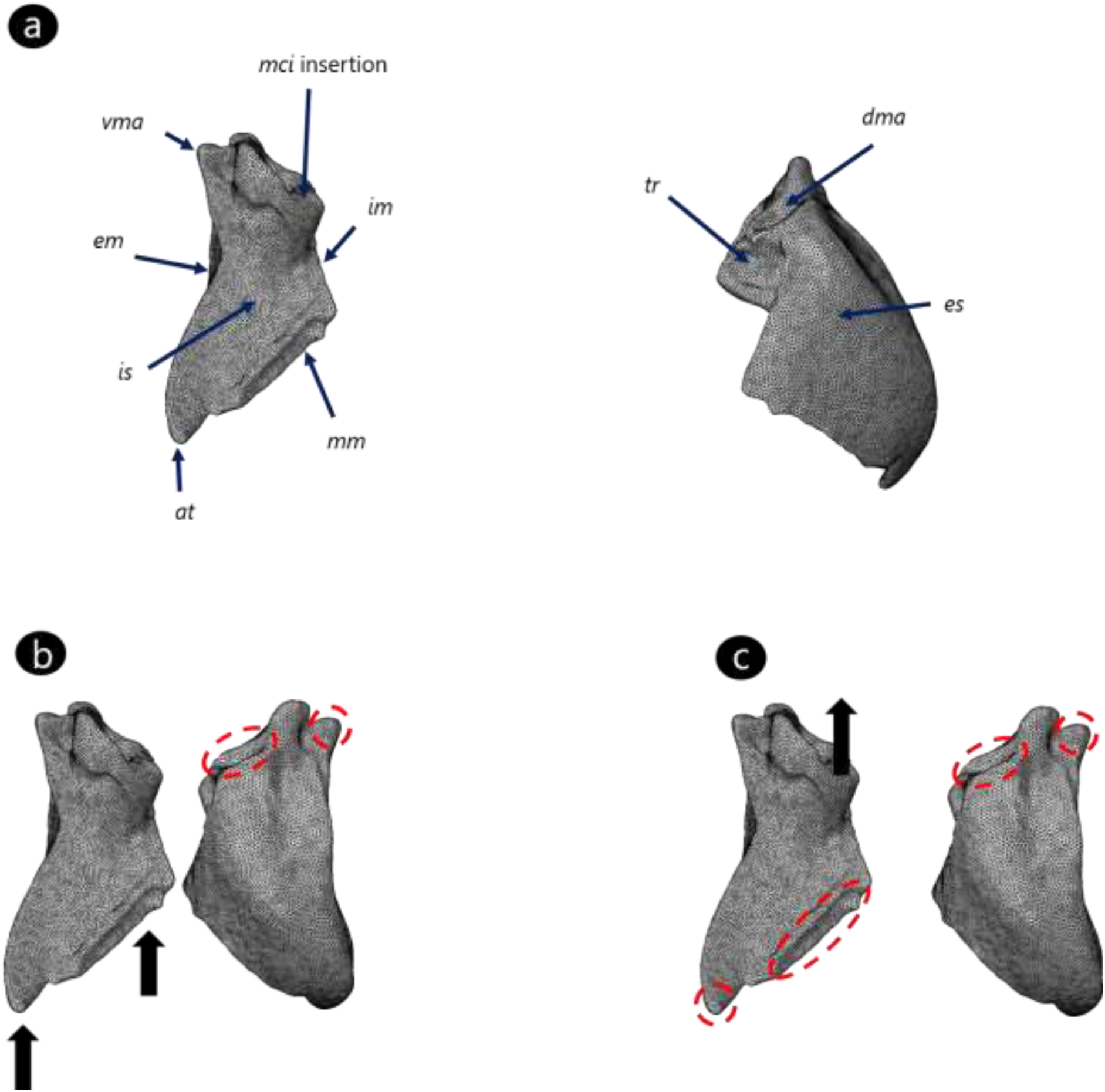
Main mandibular regions considered to discuss stress dissipation patterns (a); Loaded and constrained regions in strike (b) and pressure (b) biting simulations. In b and c, arrows indicate the direction and region of load, and dashed lines enclose the constrained regions for each simulation. *at*: apical tooth; *dma*: dorsal mandibular articulation; *em*: external margin; *es*: external surface; *im*: internal margin; *is*: internal surface; *mci*: muscle craniomandibularis internus; *mm*: masticatory margin; *tr*: trulleum; *vma*: ventral mandibular articulation.

## Results

### FEA simulations

Stress distribution results are shown in Figure 3. Given that the volume of each model varies, and that we use idealized loads and material properties, we chose not to interpret absolute stress values. Rather, we will focus on qualitative differences among simulations by rescaling the stress ranges based on a reference model to facilitate comparisons between species, subcastes, and biting scenarios. Therefore, relative differences in stress distribution between simulations indicate mandibular biomechanical distinctions to assimilate loading conditions.

**Fig. 3:**
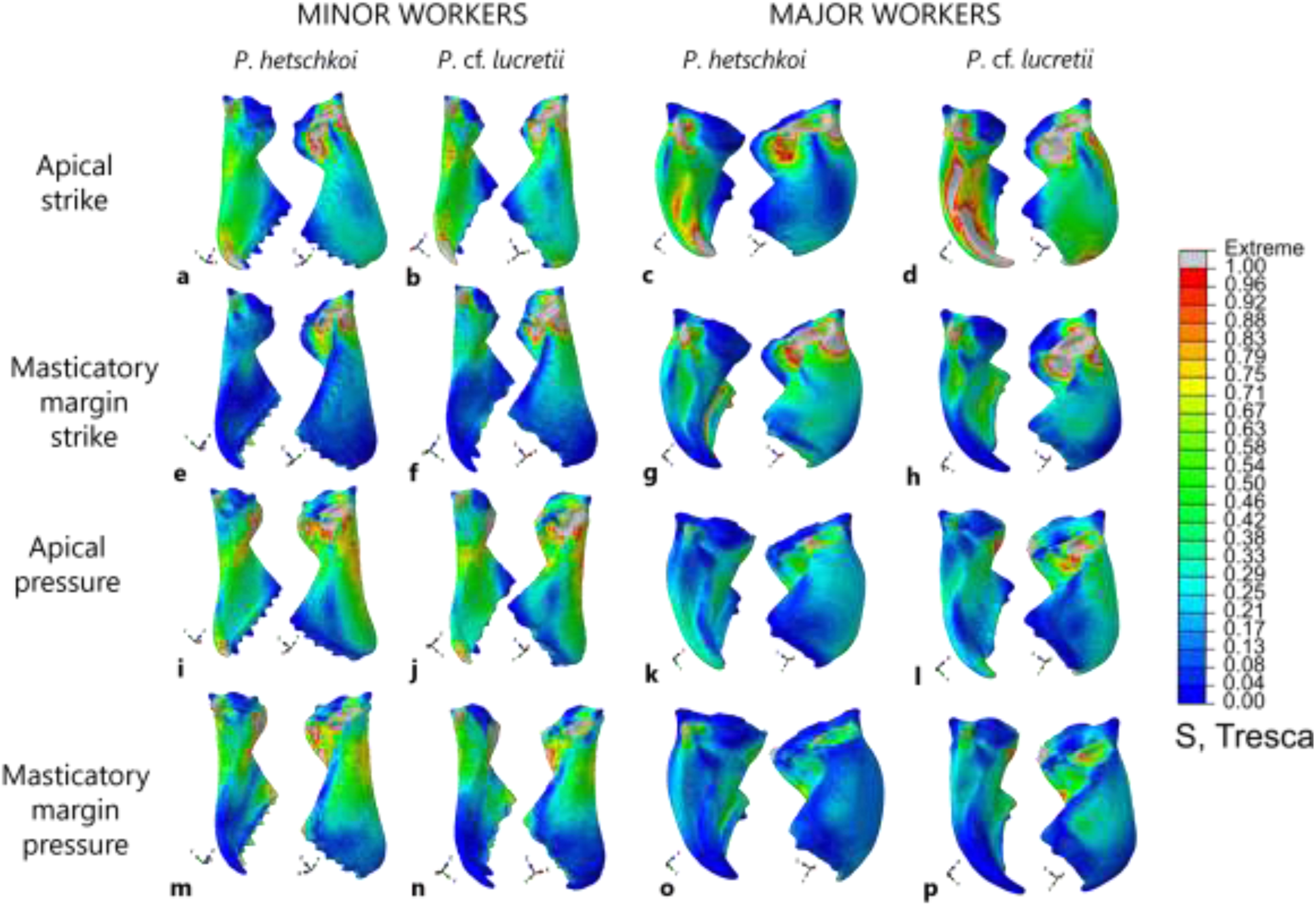
Tresca stress results (rescaled to range between 0-1) for the four biting scenarios (rows), from minors and majors of both *Pheidole* species (columns). Each letter depicts a distinct simulation. Color represents a proportional value of stress in relation to the maximum value considered for each simulation, indicated as 1.00, and grey represents extremes values above the maximum considered.

### Major worker mandibles

When displacement restrictions were applied on the mandibular joints, those regions expectedly showed high-stress levels, but stresses had to spread to other regions to be effectively absorbed. Starting from the dorsal mandibular articulation (*dma*), stresses dissipate mainly along the mandible external surface and trulleum (Fig. 3c, d, g, h, k, l, o, p). Indeed, the trulleum was important to concentrate stresses coming from the *dma* in all simulations. Stresses from the ventral mandibular articulation (*vma*) spread mainly along the external margin and through its surroundings along internal and external surfaces (Fig. 3c, d, g, h, k, l, o, p). Contrasting different biting scenarios, higher stresses are found when only the apical tooth is employed (Fig. 3c, d, k, l). This result indicates that ants face marked mechanical restrictions when they only use the apical tooth. Pressure scenarios generated higher stresses around the basal region of the internal surface (Fig. 3k, l, o, p), whereas strike scenarios concentrated more stress near the masticatory margin, an expected consequence of load application (Fig. 3c, d, g, h). However, the key aspect relating different biting scenarios are the higher stress levels in *dma* and *vma* in the strike (Fig. 3c, d, g, h) versus pressure simulations (Fig. 3k, l, o, p), which indicates that strike causes higher mechanical demands in the mandibular joints than pressure.

The main aspect that influences stress dissipation differences between species is the mandible internal surface concavity. *Pheidole hetschkoi* has a deeper concavity near the masticatory margin, which acts as an important stress concentrator, mainly in strike scenarios on the masticatory margin (Fig. 2g). While *P*. cf. *lucretii* also shows stress concentration at the same region in this biting scenario, those stresses spread more extensively along the internal surface (Fig. 3h), which suggests that its concavity is shallow and does not act as a stress concentrator. The external surface curvature also differs between species, but there are no substantial differences in terms of stress dissipation pattern (Fig. 3c, d, g, h, k, l, o, p). The dissipation through the external surface is more restricted to the articulations surroundings, given the robustness of the mandibular base, which could explain why there is not a conspicuous effect of the external surface curvature in the stress dissipation pattern between species. Stresses were proportionally higher in the *P*. cf. *lucretii* mandible, through most mandibular regions and all biting scenarios, but the differences are more striking in pressure scenarios (Fig. 3l, p).

### Minor worker mandibles

There is a distinguished stress concentration around the more constricted region of the mandibular internal surface, a trend that occurs mainly in strike simulations, especially when the load was applied on the masticatory margin (Fig. 3e, f). This constriction acts as a stress concentrator in minors due to their slender mandibles in comparison to majors. When the results of different species are compared, *P*. cf. *lucretii* simulations show proportionately higher stresses than *P. hetschkoi* in general (Fig. 3b, f, j, n), contrary to the expectation that minors mandibles would not differ in mechanical performance. The overall lower stress levels found in masticatory margin strike simulations of *P. hetschkoi* minors seems to reflect the presence of well-developed teeth along its masticatory margin. It is noticeable that the masticatory margin teeth absorb great levels of stress (Fig. 3e), so that their absence leads to higher stress levels along the mandible surfaces in strike simulations of *P*. cf. *lucretii* minor, as well as in majors of both species. The higher stresses along the internal surface in *P*. cf. *lucretii* minor mandible, compared to *P. hetschkoi* minor mandible, draw attention to the mechanical limitations associated with worn mandibles, as is the case of the *P*. cf. *lucretii* minor mandible modeled, which can lead to behavioral switches in task performance along the worker lifetime. Regarding the biting scenarios, pressure in minors result in higher stresses on internal and external surfaces of both species when compared to majors (Fig. 3i, j, m, n). As occurred in pressure scenarios for majors, stresses along the internal surface concentrate near the mandible base, where the load was applied. However, in minors, the mandible base is slender, which can explain why the mandibular surfaces in minors are proportionally more stressed in pressure than in strike simulations.

## Discussion

The use of biomechanical simulations to explore the relationships between form and function in insects is a recent endeavor. To date, FEA simulations have been used to understand biomechanical consequences of male stag beetle mandibles during agonistic interactions (Goyens et al., 2014; 2015; 2016), as well as the morphological evolution of Anisoptera (Odonata) mandibles (Blanke et al., 2017). More recently, it was demonstrated how mandibles of the trap-jaw ant *Mystrium camillae* are morphologically adapted to deal with the loadings arising from the power amplification mechanism of its closing movement (Larabee et al., 2018). Here we apply FEA in mandibles of *Pheidole* workers to simulate different biting scenarios and investigate how morphological differences in mandible morphology reflects their responses to those bite loading demands.

Our results demonstrate how mandible morphology of dimorphic workers can be optimized for particular tasks and draws attention to the role of specific mandibular regions or structures to deal with the stresses generated by their bite. Ant workers have a typical pterygote triangular mandible (Snodgrass, 1935), which can be divided into two components, a basal thick stem, and a distal blade (Richter et al., 2020). Our results indicate that this distribution of cuticular material in the mandible may conform to the high loading demands experienced by the mandibular articulations with the head. Most of the stresses generated on the apical tooth dissipate along the external margin towards the mandibular base, in both species and subcastes, avoiding the spread of considerable stresses through the more delicate mandibular surfaces. In masticatory margin stress simulations, the presence of well-developed teeth results in stresses being concentrated on the teeth instead of spreading through the internal surface. Majors of *Pheidole*, in which the masticatory margin is toothless, show a higher degree of concavity on their internal surfaces, especially in *P. hetschkoi*, which helps to concentrate stresses near the more robust masticatory margin instead of spreading through the internal surface. Although alleviating the level of stress in the mandibular articulations, such stress concentration can be harmful in cases in which the structure is submitted to cycles of loading, leading to structural fail due to material fatigue (Dirks et al., 2013).

An important aspect of *Pheidole* mandibular morphology with respect to bite mechanics is the role of the trulleum on stress concentration. The trulleum is a concavity near the *dma* present only in some myrmicine ants (Richter et al., 2019). The function of the trulleum was hitherto unknown. Here we demonstrate for the first time the importance of the trulleum to concentrate stresses coming from the *dma*, avoiding the spread of stresses through the more delicate mandibular surfaces. This is an interesting discovery, given that the *dma* seems to concentrate higher stresses in general than the *vma*. Given the trulleum clear functional role here outlined, it would be interesting to investigate the biomechanical responses of mandibles that lack the development of the trulleum, to understand how stresses dissipate from *dma* without this important stress concentrator, especially in ant species with similar loading demands as *Pheidole* mandibles.

Our results also underscore how the more robust major mandibles are better suited to deal with pressure biting than minors slender mandibles, which show higher performance in strike scenarios. As expected, these results agree with the specialized roles played by major workers in the colony. The behavioral repertoire of major workers is particularly limited, being frequently restricted to defense and/or food processing (Wilson 1984; 2003). Indeed, when minors are experimentally removed from the colony, major workers take over many of their tasks, although with decreased efficiency (Wilson 1984; Mertl & Traniello, 2009). Major mandibles meet the demands to deal with the processing of hard food items through pressure, with their toothless masticatory margin spreading bite forces evenly around the food item. Seed consumption is considered an important aspect in the evolution of several myrmicine genera, such as *Pheidole, Pogonomyrmex*, and *Solenopsis* (Ferster et al., 2006; Moreau 2008). However, the influence of granivory on morphological evolution, especially regarding the dimorphism in the *Pheidole* worker caste, is still poorly known (Holley et al., 2016). Here, we demonstrate for the first time how ant mandible morphology can be tuned to deal with the mechanical demands of processing hard food items, such as seeds and arthropod cuticles, through major mandibles better performance in pressure biting conditions. Also, mandibles of *P. hetschkoi* majors show an even better performance in pressure bite than *P*. cf. *lucretii*, suggesting that majors of *P. hetschkoi* can deal better with harder food items than *P*. cf. *lucretii*. These results may lead to the possibility of food partitioning among *Pheidole* coexisting species, and agrees with the habit of seed consumption by *P. hetschkoi*, which demands higher bite forces and consequently leads to higher stress levels on the mandibles.

Although minor workers of *Pheidole* typically show well-developed teeth in their masticatory margin, the particular specimen of *P*. cf. *lucretii* included in our study showed high levels of teeth wear, allowing us to assess the consequences of teeth wear on bite loadings. Teeth concentrate the forces generated by the masticatory muscles on smaller areas, improving the initiation of the fracture (Clissold 2007). The importance of teeth to task efficiency was demonstrated for leaf-cutting ants, where workers specialized to cut leaves switch to carrying them once their teeth are worn to a certain degree, reducing their cutting efficiency (Schofield et al., 2011). In *Pheidole*, minors perform a wide range of tasks in the colony (Wilson 1984; 2003), but information on the role of teeth wear in minor task switching is scarce. Here we demonstrate the possible mechanical consequences of teeth wear in ant mandibles, comparing the relative amount of stress generated during masticatory margin strike simulations in *P. hetschkoi* and *P*. cf. *lucretii* minors. Our results indicated that *P*. cf. *lucretii* has relatively higher stresses than *P. hetschkoi*, mainly along its mandible internal surface, which drives to higher stresses at the mandibular articulations with the head. Further studies in task allocation and mandible morphology in dimorphic ants colonies can address if teeth wear generates task switch, and biomechanical studies can reveal how teeth wear reduces task efficiency (Schofield et al., 2011).

The morphological evolution of *Pheidole* might be strongly driven by differences in size (Pie & Traniello, 2007), which tends to evolve at higher rates than shape (Pie & Tschá, 2013; Economo et al., 2015a; Friedman et al., 2019). More recently, studies applying geometric morphometric approaches validated the prominence of size to explain the morphological disparity in the genus but also pointed to different evolutionary rates and levels of integration between head and mesosoma shape and size (Friedman et al., 2019; 2020). *Pheidole* morphological diversification seems to be very constrained (Pie & Traniello, 2007), in contrast to their ecological disparity (Economo et al., 2015a; 2015b), as reflected in the widespread distribution of genus throughout most of the terrestrial ecosystems (Economo et al., 2019). Field observations demonstrate that, despite the relative morphological resemblance in *Pheidole* species, they can show considerable ecological and behavioral diversity (Mertl et al., 2010; Tschá & Pie, 2019). We demonstrate that even small morphological differences in mandibles shape between species can lead to biomechanical specialization, mainly in terms of the food processing capacity of majors. This biomechanical specialization can expand the diet range of species and lead to food partitioning (Blüthgen et al., 2003; Rosumek 2017), decreasing the degree of competition and allowing for species coexistence (Blüthgen & Feeldar, 2010).

Our results provide a biomechanical basis to understand how mandible morphological evolution can improve task specialization in polymorphic ants. Morphological polymorphism in the worker caste can expand the range of prey items that a species is able to handle, as demonstrated for some species of the army ant genus *Eciton* (Powell & Franks, 2005; 2006). In the highly polymorphic genus *Cephalotes*, which is *Pheidole*’s sister lineage, some workers are specialized to plug the nest entrances with their heads to protect the colony against invasion (Powell 2008). In some fire ants such as *Solenopsis geminate* (Fabricius), the degree of worker polymorphism is associated to higher levels of division of labor, with major workers being specialized to seed milling (Wilson 1978; Ferster et al., 2006). Division of labor in leaf-cutting ants is associated with morphological distinctions among worker mandibles, as demonstrated for the polymorphic genus *Atta* (Silva et al., 2016). Therefore, the role of worker polymorphism for division of labor in ants is well established (Wills et al., 2018), but by applying biomechanical approaches we can advance our understanding about the functional role of morphological disparity, as we demonstrated here for *Pheidole* workers. In this sense, ant polymorphic lineages are ideal models to investigate form-function relationships, and the morphological differentiation of mandibles should be studied in detail, given the importance of this structure to worker interactions with the environment.

## Acknowledgments

The authors acknowledge Dr. Emily Rayfield, Dr. Mauricio Moura and Dr. Roberto Keller for valuable comments on our manuscript. Thanks to the CAPES Foundation for the support provided to CLK (doctorate scholarship - 001) and ACF (PDSE grant 88881.189085/2018-01). E.P.E. was supported by subsidy funding to OIST. The authors thank the OIST Imaging section for access to the CT scanner.

## Supporting Information

**Table SI1:** Characteristics of each finite element mesh.

|  | Specimen | Mesh volume (mm<sup>3</sup>) | Element edge length (mm) | Number of elements |
| --- | --- | --- | --- | --- |
| Major | *P. hetschkoi* | 0.0741 | 0.023 | 449488 |
|  | *P. cf. lucretii* | 0.0183 | 0.023 | 278634 |
| Minor | *P. hetschkoi* | 0.00469 | 0.0035 | 881691 |
|  | *P. cf. lucretii* | 0.00191 | 0.0035 | 392790 |

## Notes

### Competing Interest Statement

The authors have declared no competing interest.

## References

Azevedo AF. 2003 Método dos Elementos Finitos. Porto: Faculdade de Engenharia da Universidade do Porto.

Blanke A, Schmitz H, Patera A, Dutel H, Fagan MJ. 2017 Form–function relationships in dragonfly mandibles under an evolutionary perspective. J. R. Soc. Interface 14, 20161038. (doi:10.1098/rsif.2016.1038)

Blüthgen N, Feldhaar H. 2010 Food and shelter: how resources influence ant ecology. In Ant Ecology, pp. 115–136. New York: Oxford University Press.

Blüthgen N, Gebauer G, Fiedler K. 2003 Disentangling a rainforest food web using stable isotopes: dietary diversity in a species-rich ant community. Oecologia 137, 426–435. (doi:10.1007/s00442-003-1347-8)

Bolton B. An online catalog of the ants of the world. 2020. antcat.org (last access in 03/SEP/2020).

Brito TO, Elzubair A, Araújo LS, Camargo SA de S, Souza JLP, Almeida LH. 2017 Characterization of the Mandible Atta Laevigata and the Bioinspiration for the Development of a Biomimetic Surgical Clamp. Mat. Res. 20, 1525–1533. (doi:10.1590/1980-5373-mr-2016-1137)

Cignoni P, Callieri M, Corsini M, Dellepiane M, Ganovelli F, Ranzuglia G. 2008 Meshlab: an open-source mesh processing tool. In Eurographics Italian chapter conference, pp. 129–136.

Clissold FJ. 2007 The biomechanics of chewing and plant fracture: mechanisms and implications. Adv In Insect Phys 34, 317–372.

Dirks J-H, Parle E, Taylor D. 2013 Fatigue of insect cuticle. Journal of Experimental Biology 216, 1924–1927. (doi:10.1242/jeb.083824)

Economo EP, Huang J-P, Fischer G, Sarnat EM, Narula N, Janda M, Guénard B, Longino JT, Knowles LL. 2019 Evolution of the latitudinal diversity gradient in the hyperdiverse ant genus Pheidole. Global Ecol Biogeogr 28, 456–470. (doi:10.1111/geb.12867)

Economo EP, Klimov P, Sarnat EM, Guénard B, Weiser MD, Lecroq B, Knowles LL. 2015a Global phylogenetic structure of the hyperdiverse ant genus Pheidole reveals the repeated evolution of macroecological patterns. Proc. R. Soc. B 282, 20141416. (doi:10.1098/rspb.2014.1416)

Economo EP et al. 2015b Breaking out of biogeographical modules: range expansion and taxon cycles in the hyperdiverse ant genus Pheidole. J. Biogeogr. 42, 2289–2301. (doi:10.1111/jbi.12592)

Evans AR, Sanson GD. 2005 Biomechanical properties of insects in relation to insectivory: cuticle thickness as an indicator of insect ‘hardness’ and ‘intractability’. Aust. J. Zool. 53, 9. (doi:10.1071/ZO04018)

Ferster B, Pie MR, Traniello JFA. 2006 Morphometric variation in North American Pogonomyrmex and Solenopsis ants: caste evolution through ecological release or dietary change? Ethology Ecology & Evolution 18, 19–32. (doi:10.1080/08927014.2006.9522723)

Friedman NR, Lecroq Bennet B, Fischer G, Sarnat EM, Huang J, Knowles LLK, Economo EP. 2020 Macroevolutionary integration of phenotypes within and across ant worker castes. Ecol Evol, ece3.6623. (doi:10.1002/ece3.6623)

Friedman NR, Remeš V, Economo EP. 2019 A morphological integration perspective on the evolution of dimorphism among sexes and social insect castes. Integr Comp Biol 59, 410–419.

Gadagkar R. 1997 The evolution of caste polymorphism in social insects: Genetic release followed by diversifying evolution. J. Genet. 76, 167–179. (doi:10.1007/BF02932215)

Goyens J, Dirckx J, Aerts P. 2015 Built to fight: variable loading conditions and stress distribution in stag beetle jaws. Bioinspir. Biomim. 10, 046006. (doi:10.1088/1748-3190/10/4/046006)

Goyens J, Dirckx J, Aerts P. 2016 Jaw morphology and fighting forces in stag beetles. J Exp Biol 219, 2955–2961. (doi:10.1242/jeb.141614)

Goyens J, Soons J, Aerts P, Dirckx J. 2014 Finite-element modelling reveals force modulation of jaw adductors in stag beetles. J. R. Soc. Interface 11, 20140908. (doi:10.1098/rsif.2014.0908)

Gronenberg W, Paul J, Just S, Hölldobler B. 1997 Mandible muscle fibers in ants: fast or powerful? Cell and Tissue Research 289, 347–361. (doi:10.1007/s004410050882)

Hao W, Yao G, Zhang X, Zhang D. 2018 Kinematics and Mechanics analysis of trap-jaw ant Odontomachus monticola. J. Phys.: Conf. Ser. 986, 012029. (doi:10.1088/1742-6596/986/1/012029)

Hölldobler B, Wilson EO. 1990 The ants. Cambridge, Mass: Belknap Press of Harvard University Press.

Holley J-AC, Moreau CS, Laird JG, Suarez AV. 2016 Subcaste-specific evolution of head size in the ant genus Pheidole. Biol. J. Linn. Soc. 118, 472–485. (doi:10.1111/bij.12769)

Huang MH, Wheeler DE, Fjerdingstad EJ. 2013 Mating system evolution and worker caste diversity in Pheidole ants. Mol Ecol 22, 1998–2010. (doi:10.1111/mec.12218)

Larabee FJ, Smith AA, Suarez AV. 2018 Snap-jaw morphology is specialized for high-speed power amplification in the Dracula ant, Mystrium camillae. R. Soc. open sci. 5, 181447. (doi:10.1098/rsos.181447)

Lillico-Ouachour A, Metscher B, Kaji T, Abouheif E. 2018 Internal head morphology of minor workers and soldiers in the hyperdiverse ant genus Pheidole. Can. J. Zool. 96, 383– 392. (doi:10.1139/cjz-2017-0209)

Marcé-Nogué J, Fortuny J, Gil L, Sánchez M. 2015 Improving mesh generation in finite element analysis for functional morphology approaches. Spanish J. Palaeontol. 30, 117–132. (doi:10.7203/sjp.30.1.17227)

Mertl AL, Sorenson MD, Traniello JFA. 2010 Community-level interactions and functional ecology of major workers in the hyperdiverse ground-foraging Pheidole (Hymenoptera, Formicidae) of Amazonian Ecuador. Insect. Soc. 57, 441–452. (doi:10.1007/s00040-010-0102-5)

Mertl AL, Traniello JFA. 2009 Behavioral evolution in the major worker subcaste of twig-nesting Pheidole (Hymenoptera: Formicidae): does morphological specialization influence task plasticity? Behav Ecol Sociobiol 63, 1411–1426. (doi:10.1007/s00265-009-0797-3)

Moreau CS. 2008 Unraveling the evolutionary history of the hyperdiverse ant genus Pheidole (Hymenoptera: Formicidae). Molecular Phylogenetics and Evolution 48, 224–239. (doi:10.1016/j.ympev.2008.02.020)

Muscedere ML, Traniello JFA, Gronenberg W. 2011 Coming of age in an ant colony: cephalic muscle maturation accompanies behavioral development in Pheidole dentata. Naturwissenschaften 98, 783–793. (doi:10.1007/s00114-011-0828-6)

Neville AC. 1975 General Structure of Integument. In Biology of the Arthropod Cuticle, pp. 7–70. Berlin: Springer-Verlag.

Oliver WC, Pharr GM. 1992 An improved technique for determining hardness and elastic modulus using load and displacement sensing indentation experiments. J. Mater. Res. 7, 1564–1583. (doi:10.1557/JMR.1992.1564)

Oster GF, Wilson EO 1978 Caste and Ecology in the Social Insects. New Jersey: Princeton University Press.

Özkaya N, Leger D, Goldsheyder D, Nordin M. 2017 Multiaxial Deformations and Stress Analyses. In Fundamentals of Biomechanics: Equilibrium, Motion, and Deformation, pp. 317–360. New York: Springer.

Paul J. 2001 Mandible movements in ants. Comparative Biochemistry and Physiology Part A: Molecular & Integrative Physiology 131, 7–20. (doi:10.1016/S1095-6433(01)00458-5)

Paul J, Gronenberg W. 1999 Optimizing force and velocity: mandible muscle fibre attachments in ants. Journal of Experimental Biology 202, 797–808.

Pie MR, Traniello JFA. 2007 Morphological evolution in a hyperdiverse clade: the ant genus Pheidole. J Zoology 271, 99–109. (doi:10.1111/j.1469-7998.2006.00239.x)

Pie MR, Tschá MK. 2013 Size and shape in the evolution of ant worker morphology. PeerJ 1, e205. (doi:10.7717/peerj.205)

Powell S, Franks NR. 2005 Caste evolution and ecology: a special worker for novel prey. Proc. R. Soc. B 272, 2173–2180. (doi:10.1098/rspb.2005.3196)

Powell S. 2008 Ecological specialization and the evolution of a specialized caste in Cephalotes ants. Functional Ecology 22, 902–911. (doi:10.1111/j.1365-2435.2008.01436.x)

Powell S, Franks NR. 2006 Ecology and the evolution of worker morphological diversity: a comparative analysis with Eciton army ants. Funct Ecology 20, 1105–1114. (doi:10.1111/j.1365-2435.2006.01184.x)

Powell S. 2009 How ecology shapes caste evolution: linking resource use, morphology, performance and fitness in a superorganism. Journal of Evolutionary Biology 22, 1004–1013. (doi:10.1111/j.1420-9101.2009.01710.x)

Rajabi H, Ghoroubi N, Darvizeh A, Dirks J-H, Appel E, Gorb SN. 2015 A comparative study of the effects of vein-joints on the mechanical behaviour of insect wings: I. Single joints. Bioinspir. Biomim. 10, 056003. (doi:10.1088/1748-3190/10/5/056003)

Rajabi H, Ghoroubi N, Darvizeh A, Appel E, Gorb SN. 2016 Effects of multiple vein microjoints on the mechanical behaviour of dragonfly wings: numerical modelling. R. Soc. open sci. 3, 150610. (doi:10.1098/rsos.150610)

Rayfield EJ. 2007 Finite Element Analysis and Understanding the Biomechanics and Evolution of Living and Fossil Organisms. Annu. Rev. Earth Planet. Sci. 35, 541–576. (doi:10.1146/annurev.earth.35.031306.140104)

Richter A, Hita Garcia F, Keller RA, Billen J, Economo EP, Beutel RG. 2020 Comparative analysis of worker head anatomy of Formica and Brachyponera (Hymenoptera: Formicidae). Arthropod Systematics & Phylogeny 78, 133–170. (doi:10.26049/ASP78-1-2020-06)

Richter A, Keller RA, Rosumek FB, Economo EP, Hita Garcia F, Beutel RG. 2019 The cephalic anatomy of workers of the ant species Wasmannia affinis (Formicidae, Hymenoptera, Insecta) and its evolutionary implications. Arthropod Structure & Development 49, 26–49. (doi:10.1016/j.asd.2019.02.002)

Rosumek FB. 2017 Natural History of Ants: What We (do not) Know About Trophic and Temporal Niches of Neotropical Species. Sociobiology 64, 244. (doi:10.13102/sociobiology.v64i3.1623)

Sarnat EM, Friedman NR, Fischer G, Lecroq-Bennet B, Economo EP. 2017 Rise of the spiny ants: diversification, ecology and function of extreme traits in the hyperdiverse genus Pheidole (Hymenoptera: Formicidae). Biological Journal of the Linnean Society 122, 514–538. (doi:10.1093/biolinnean/blx081)

Schofield RMS, Emmett KD, Niedbala JC, Nesson MH. 2011 Leaf-cutter ants with worn mandibles cut half as fast, spend twice the energy, and tend to carry instead of cut. Behav Ecol Sociobiol 65, 969–982. (doi:10.1007/s00265-010-1098-6)

Silva LC, Camargo RS, Lopes JFS, Forti LC. 2016 Mandibles of Leaf-Cutting Ants: Morphology Related to Food Preference. Sociobiology 63, 881. (doi:10.13102/sociobiology.v63i3.1014)

Sirohi J, Chopra I. 2000 Fundamental Understanding of Piezoelectric Strain Sensors. Journal of Intelligent Material Systems and Structures 11, 246–257. (doi:10.1106/8BFB-GC8P-XQ47-YCQ0)

Snodgrass RE, 1935 Principles of Insect Morphology. New York: Cornell University Press.

Sun J, Tong J, Ma Y. 2008 Nanomechanical Behaviours of Cuticle of Three Kinds of Beetle. Journal of Bionic Engineering 5, 152–157. (doi:10.1016/S1672-6529(08)60087-6)

Tschá MK, Pie MR. 2019 Correlates of ecological dominance within Pheidole ants (Hymenoptera: Formicidae): Correlates of ecological dominance in ants. Ecol Entomol 44, 163–171. (doi:10.1111/een.12685)

Vincent JFV, Wegst UGK. 2004 Design and mechanical properties of insect cuticle. Arthropod Structure & Development 33, 187–199. (doi:10.1016/j.asd.2004.05.006)

Yushkevich PA, Piven J, Hazlett HC, Smith RG, Ho S, Gee JC, Gerig G. 2006 User-guided 3D active contour segmentation of anatomical structures: Significantly improved efficiency and reliability. NeuroImage 31, 1116–1128. (doi:10.1016/j.neuroimage.2006.01.015)

Wheeler DE. 1991 The Developmental Basis of Worker Caste Polymorphism in Ants. The American Naturalist 138, 1218–1238. (doi:10.1086/285279)

Wheeler WM. 1910. Ants: their structure, development and behavior. New York: Columbia University Press.

Wills BD, Powell S, Rivera MD, Suarez AV. 2018 Correlates and Consequences of Worker Polymorphism in Ants. Annu. Rev. Entomol. 63, 575–598. (doi:10.1146/annurev-ento-020117-043357)

Wilson EO. 1978 Division of labor in fire ants based on physical castes (Hymenoptera: Formicidae: Solenopsis). Journal of the Kansas Entomological Society 51, 615–636.

Wilson EO. 2003 Pheidole in the New World: a dominant, hyperdiverse ant genus. Cambridge, Mass: Harvard University Press.

Wilson EO. 1971 The insect societies. Cambridge, EUA: Harvard University Press.

Wilson EO. 1953 The Origin and Evolution of Polymorphism in Ants. The Quarterly Review of Biology 28, 136–156. (doi:10.1086/399512)

Wilson EO. 1984 The relation between caste ratios and division of labor in the ant genus Pheidole (Hymenoptera: Formicidae). Behav Ecol Sociobiol 16, 89–98. (doi:10.1007/BF00293108)

